# DE-NOVO HEMATOPOIESIS FROM THE FETAL LUNG

**DOI:** 10.1101/2022.02.14.480402

**Authors:** Anthony K. Yeung, Carlos Villacorta-Martin, Jonathan Lindstrom-Vautrin, Anna C. Belkina, Kim Vanuytsel, Todd W. Dowrey, Alexandra B. Ysasi, Vladimir Vrbanac, Gustavo Mostoslavsky, Alejandro B. Balazs, George J. Murphy

## Abstract

Hemogenic endothelial cells (HECs) are specialized cells that undergo endothelial to hematopoietic transition (EHT) to give rise to hematopoietic progenitors. Though not defined as a hematopoietic organ, the lung houses many resident hematopoietic cells, aids in platelet biogenesis, and is a reservoir for hematopoietic stem and progenitor cells (HSPCs), but lung HECs have never been described. Using explant cultures of murine and human fetal lungs, we demonstrate that the fetal lung is a source of HECs that have the functional capacity to undergo EHT to produce de-novo HSPCs. Flow cytometric and functional assessment of fetal lung explants showed the production of HSPCs that expressed key EHT and pre-HSPC markers. scRNA-Seq and small molecule modulation demonstrated that fetal lung EHT is reliant on canonical EHT signaling pathways. These findings suggest that functional HECs are present in the fetal lung, thus establishing this location as a potential extramedullary site of de-novo hematopoiesis.

## INTRODUCTION

Hematopoietic stem cells (HSCs) originate from a rare sub-population of arterial endothelial cells known as HECs. Making up just 1-3% of the total endothelial cell population in the AGM, HECs are commonly thought to be confined to a small window of gestation between embryonic days 8 to 11 (E8-11) in mice, and E27-40 in humans^1,2^. Within this time window, HECs can also be found in other hematopoietic organs and vessels including the yolk sac, placenta, and vitelline and umbilical arteries^3–5,6(p1),7,8^. Some studies suggest that HECs may not be restricted to these hematopoietic organs and developmental window. Work from others suggests that functional HECs may also be found in the embryonic head around E10-E11 as well as the perinatal chicken and murine bone marrow (BM)^9,10^. Collectively, these studies demonstrate that our spatiotemporal understanding of HECs remains limited and raises questions about the hematopoietic potential of other organs.

Recent studies have highlighted the presence and importance of various resident blood cell populations in the lung^11^. In particular, the lung was demonstrated to be a reservoir of HSPCs, but the origins of this population of progenitors remains unknown^12–14^. *In-utero* mechanical stimuli have been shown to be an important environmental cue for both HEC and pulmonary development. More specifically, cyclic stretch induced activation of Yes Activated Protein (YAP) signaling in the AGM promotes EHT^15^. Interestingly, rhythmic breathing movements begin occurring around E16 in fetal mice, and mechanical stimuli has a significant impact on fetal airway and alveolar epithelial development^16–18^. Beyond mechanical signaling, cell-cell extrinsic signaling within the AGM niche is critical for EHT. The ventral wall of the dorsal aorta is the most common site of EHT due to its proximity to the underlying mesenchyme which modulates EHT via Notch, BMP4, SHH, and Wnt pathways^3,19,20^. All of these pathways are similarly very important in pulmonary fetal development^21^.

Based on these parallels, we hypothesized that the fetal lung is another potential site of hemogenic endothelium with the functional capacity to produce hematopoietic progenitors. Employing fetal lung explant cultures, we demonstrate that the murine and human fetal lung are sources of putative HECs. Multipotent, fetal lung-derived HSPCs showed the canonical features of cells produced by EHT as determined by flow cytometry, single cell transcriptomics, functional assays and immunofluorescent histology. These findings highlight the fetal lung as another potential site of de-novo hematopoiesis, suggesting that the lung may have a greater role in instructing tissue specific hematopoiesis and/or overall hematopoietic development.

## MATERIALS AND METHODS

### Ethics Statement

All animal housing and experimental procedures were approved by the Boston University School of Medicine Institutional Animal Care and Use Committee (BUSM IACUC). Work involving human tissue samples was approved by Partners Human Research Committee (Protocol #2016P001106).

### Tissue isolation and processing

#### Mouse

E17 timed-pregnant C57/BL6 mice were purchased from Jackson Laboratories. The fetal liver and lung were isolated from surrounding tissue by blunt dissection and set aside in a solution of 10% characterized fetal bovine serum in Hanks’ balanced salt solution (HBSS, Gibco). Using a 5mL syringe fitted with a 16-gauge needle, the fetal liver and fetal lung were drawn up and expelled several times to physically dissociate the tissue. Samples were subsequently placed in a digest buffer containing HBSS, 1mg/mL DNase I (Sigma), and 0.5mg/mL LiberaseTM (Sigma). This digest mixture was placed on a rocker at 37°C for 30-minutes to 1-hour. Post-digestion, lung samples were filtered and resuspended in RBC lysis for 5 min at 37°C, then washed, filtered, and resuspended in HSPC medium.

#### Human

All fetal samples were within the age range between 19-24 weeks post-conception. Lung lobes were dissected away from the main airways and minced with a scalpel before being placed in a digest buffer containing HBSS, 1mg/mL DNase I (Sigma), and 0.5mg/mL LiberaseTM (Sigma). This digest mixture was placed on a rocker at 37°C for 30-minutes to 1-hour. Post digestion, lung samples were filtered and resuspended in RBC lysis for 5 min at 37°C, then washed, filtered, and resuspended in HSPC medium.

### Explant Cultures

Isolated cells suspended in HSPC medium were plated onto Matrigel coated plates. HSPC medium was made up of StemPro-34 Serum Free Medium, 50 µg/ml ascorbic acid, 400 nM monothioglycerol, 100µg/ml Primocin, 2 mM L-glutamine, and the following human or murine growth factors: 50 ng/ml vascular endothelial growth factor A (VEGFA), 100 ng/ml basic fibroblast growth factor (bFGF), 100 ng/ml stem cell factor (SCF), 100 ng/ml FMS-related tyrosine kinase ligand (FLT3L), 100 ng/ml thrombopoietin (TPO), 100 ng/ml Interleukin-6 (IL6). On the third day of all cultures, media was aspirated, rinsed once with PBS, and fresh media was applied.

### MethoCult Colony-Forming Unit (CFU) Assay

CFU assay was performed using the murine MethoCult GF M3434 (Stem Cell Techologies) or human MethoCult H4034 Optimum (Stem Cell Techologies) kit. Procedure was performed per manufacturer’s instructions.

### Tissue Histology and Cell Imaging

A small portion of the human fetal lung was fixed in 4% paraformaldehyde for 2 hours, cryoprotected in 30% sucrose and embedded in Optimum Cutting Temperature embedding medium. 10-12µm tissue sections were made using a cryostat. Sections were rinsed with PBS prior to permeabilizing and blocking in a solution of 0.4% Triton X-100 and 10% NDS. A list of antibodies used for immunofluorescence can be found under **Supplemental Table 1**.

Imaging of cell cultures were performed using a Keyence BZ-X700 fluorescence microscope. Cytospins were stained using the Hema 3 Stat Pack (Fisher) and imaged using a Nikon Eclipse NiE.

### Flow cytometry

All staining and washing steps were done using a solution of 1% BSA in PBS, 5mM EDTA. Single cell suspensions were stained with the appropriate master mix of antibodies on ice for 30 minutes. A table of antibodies used for analysis can be found under **Supplemental Table 1**. Fc-Receptor block treatment was performed using an anti-mouse CD16/32 antibody (BioLegend, Clone: 93) or anti-human Fc receptor block (biolegend, Human TruStain FcX) for 10 minutes prior to the addition of master mix. True-Stain Monocyte Blocker (BioLegend) and Brilliant Stain Buffer Plus (BD Biosciences) were included in master mix to minimize non-specific staining. Single stain controls were processed using cells and/or beads (Invitrogen, UltraComp eBeads Plus). Flow cytometry was performed using either a BD LSRII, a Stratedigm S1000EXI or a 5-laser Cytek Aurora spectral flow cytometer. Analysis was performed using FlowJo_v10.8 (FlowJo, LLC) software.

### Single cell RNA sequencing

Adherent cells from days 4, 5 and 6 of explant cultures were released from Matrigel coated wells using Accutase. Collected cells were stained for VE-Cadherin-APC-Cy7 and resuspended in sort buffer (2% BSA in PBS) containing Calcein Blue AM (1:1000). Using a Beckman MoFlo Coulter Astrios, live-singlets and VECAD^+^ events were sorted into sort buffer (2% BSA in PBS with 5mM EDTA). Cells counts and viability after sorting were confirmed by hemocytometer using trypan blue. Suspension cells were left unsorted as VECAD expression is lost gradually as hematopoietic cells differentiate post-EHT emergence. VECAD^+^ sorted adherent cells and unsorted suspension cells were single cell captured using the 10X Genomics Chromium platform and prepared using the Single Cell 3’ v3 kit. Library preparation and sequencing was done at the Boston University Microarray and Sequencing Resource (BUMSR) Core using the Illumina NextSeq 2000 instrument.

### Bioinformatic analysis

Reads were demultiplexed and aligned to the mouse genome assembly (GRCm38, Ensembl) with the STARsolo pipeline^22^. Further analyses were done using Seurat v. 3.1.4^23^. After inspection of the quality control metrics, cells with more than 12% of mitochondrial content or less than 800 detected genes were excluded for downstream analyses. We normalized and scaled the UMI counts using the regularized negative binomial regression (SCTransform)^24^. Following the standard procedure in Seurat’s pipeline, we performed linear dimensionality reduction (PCA), and used the top 20 principal components to compute both the UMAP^25^ and the clusters (Louvain method^26^) which were computed at a range of resolutions from 1.5 to 0.05 (more to fewer clusters). Cell cycle scores and classifications were done using the Seurat method^27^. The cut-offs for independent filtering^28^ prior to differential expression testing required genes: a) being detected in at least 10% of the cells of either population and b) having a natural log fold change of at least 0.25 between populations. The tests were performed using Seurat’s wrapper for the MAST framework^29^. For a comparison on the performance of methods for single-cell differential expression see Soneson & Robinson et al.^30^.

### Statistical analysis

Significance for Enrichr based pathway enrichment was determined by Fisher exact test^31,32^. Significance for pairwise scRNA-Seq gene expression comparisons was determined using the MAST framework^29^. Significance for flow cytometry assays performed in biological triplicate was determined by student paired t-test.

## RESULTS

### Murine fetal lung explant cultures produce HSPCs

To assess the functional capacity of potential lung HECs to produce blood progenitors, cells isolated from murine E17 lungs were plated onto Matrigel coated plates in an adapted hematopoietic differentiation media, termed HSPC medium^33^. This serum-free media was formulated to support the final stages of specification of HECs into HSPCs. The primary rationale for choosing E17 was that (1) this is during the window that fetal breathing movements are occurring^16^, and (2) E17 is a timepoint that is distant from the AGM EHT window (E8-11)^1^, thus helping to minimize contamination from AGM-EHT derived progenitors.

In contrast to fetal liver explants where immediate expansion of floating cells is observed due to an expanding hematopoietic progenitor population, fetal lung explants initially developed a robust adherent layer. Discrete clusters of suspension cells are observed by day 3, which expand to robust colonies between days 4-6 (**Figure 1A**). To visually determine cell identity based on their morphology, cytospins of day 6 suspension cells were performed, which revealed cells with a progenitor-like morphology as well as various differentiated hematopoietic cells including macrophages/monocytes, neutrophils and megakaryocytes (**Figure 1B**). This diversity of hematopoietic cells suggests that hematopoietic progenitors are arising from these cultures. To examine this, suspension cells were functionally assessed for progenitor potential by MethoCult assay. Day 4-6 suspension cells showed similar potential to form all types of colony forming units (CFUs), but this potential was no longer detected at Day 8 (**Figure 1C, D**). Progenitor phenotyping was further assessed by flow cytometric assessment of the broad murine HSPC markers Lin^-^/Sca^+^/Kit^+^ (LSK) and the SLAM markers CD48 and CD150. A greater fraction of CD45^+^ hematopoietic cells isolated from the adherent cell layer were LSK-HSPCs and CD48^-^/CD150^+^ SLAM marker defined HSCs (**Figure 1E**). Collectively, these data suggest that the adherent layer of cells from fetal lung explants gave rise to HSPCs.

**Figure 1.**
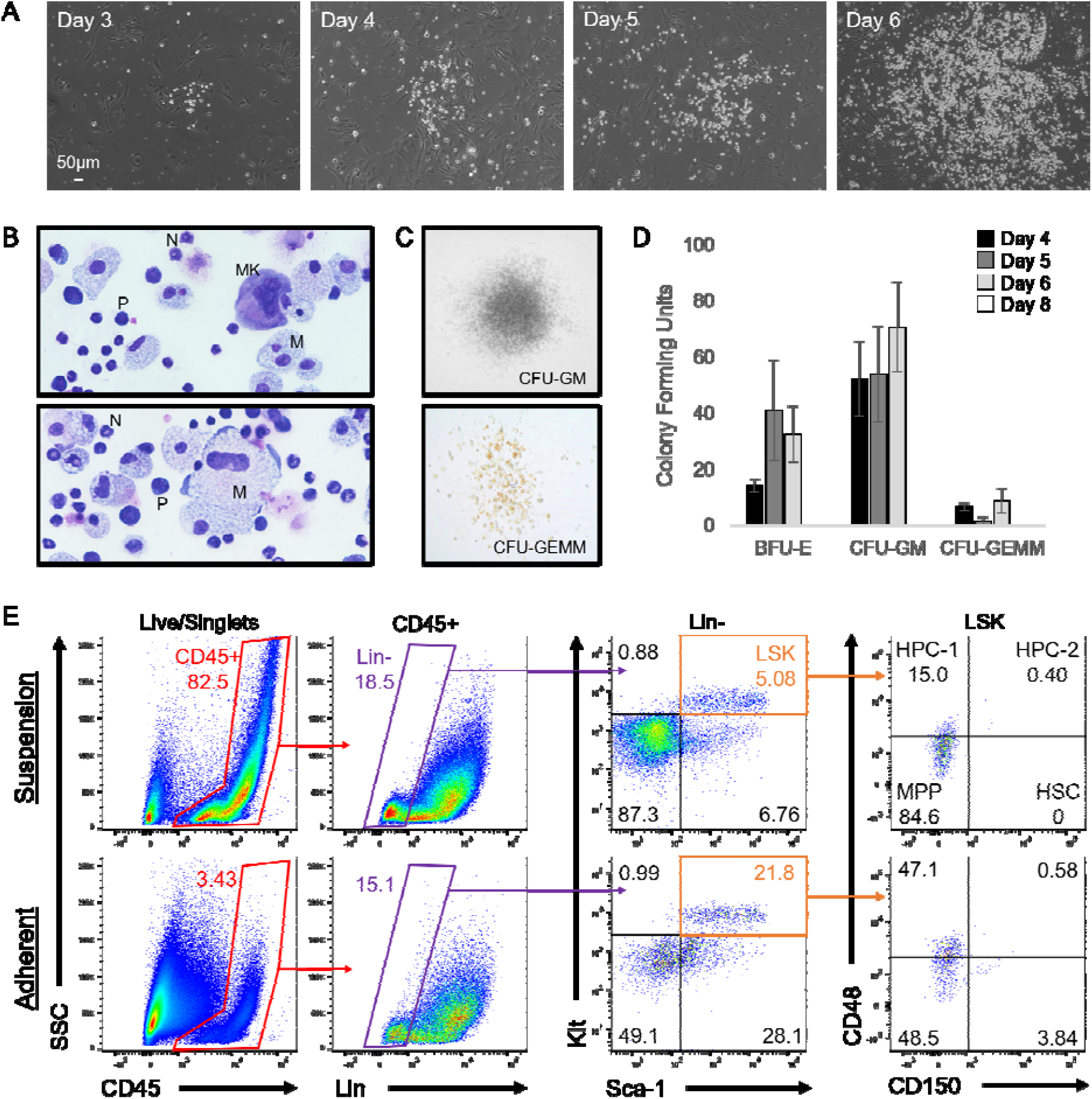
Murine fetal lung explant cultures produce HSPCs. (A) Images of a single position between days 3 to 6 of a murine fetal lung explant culture. (B) Cytospin of fetal lung derived suspension cells. Images are annotated for cell type: MK = Megakaryocyte, N = Neutrophil, M = Monocyte/Macrophage, P = Progenitor. (C) Images of representative colonies from a CFU assay. (D) Colony counts from CFU assays performed with day 4, 5, 6 and 8 suspension cells. (E) Representative plots of flow cytometric assessment of LSK and SLAM marker defined HSPC populations. Suspension represents cells that were floating in media, and Adherent represents cells that were collected post-treatment with Accutase. Error bars represent standard deviation.

### Murine fetal lung explants exhibit the dynamics of EHT

Time-lapse capture of live explant cultures from days 5-6 showed adherent cells transitioning to suspension cells (**Supplemental_Video**). These observations closely mimic AGM explant cultures and in-vitro based hematopoietic differentiations showing the transition of HECs into hematopoietic cells^34–37^. The resultant pre-HSPCs that are birthed from AGM derived HECs retain some endothelial markers and are marked by co-expression of the endothelial marker vascular endothelial cadherin (VECAD) and the broad hematopoietic marker CD45^38^. Fetal Lung explants gave rise to VECAD^+^/CD45^+^ cells, and a gradual reduction in VECAD expression coincided with the expression of the more differentiated marker CD45 (**Figure 2A**). The fraction of CD45^+^ cells that were VECAD^+^ also decreased with successive days of culture (**Figure 2B**). Time matched fetal livers, which house HSPCs but not HECs^39^, were cultured under the same conditions and did not result in the production of a population of VECAD^+^/CD45^+^ cells (**Supplemental Figure 1A**). This suggests that fetal lung derived VECAD^+^/CD45^+^ pre-HSPCs are not resultant from the expansion of pre-existing progenitors. SLAM marker defined HSCs were predominantly present amongst the VECAD^+^ suspension cells (**Figure 2C**). With time, a gradual reduction in the fraction of VECAD^+^ HSCs was observed, which coincided with the expansion of the more lineage restricted hematopoietic progenitor cell 1 (HPC-1) and HPC-2 populations.

**Figure 2.**
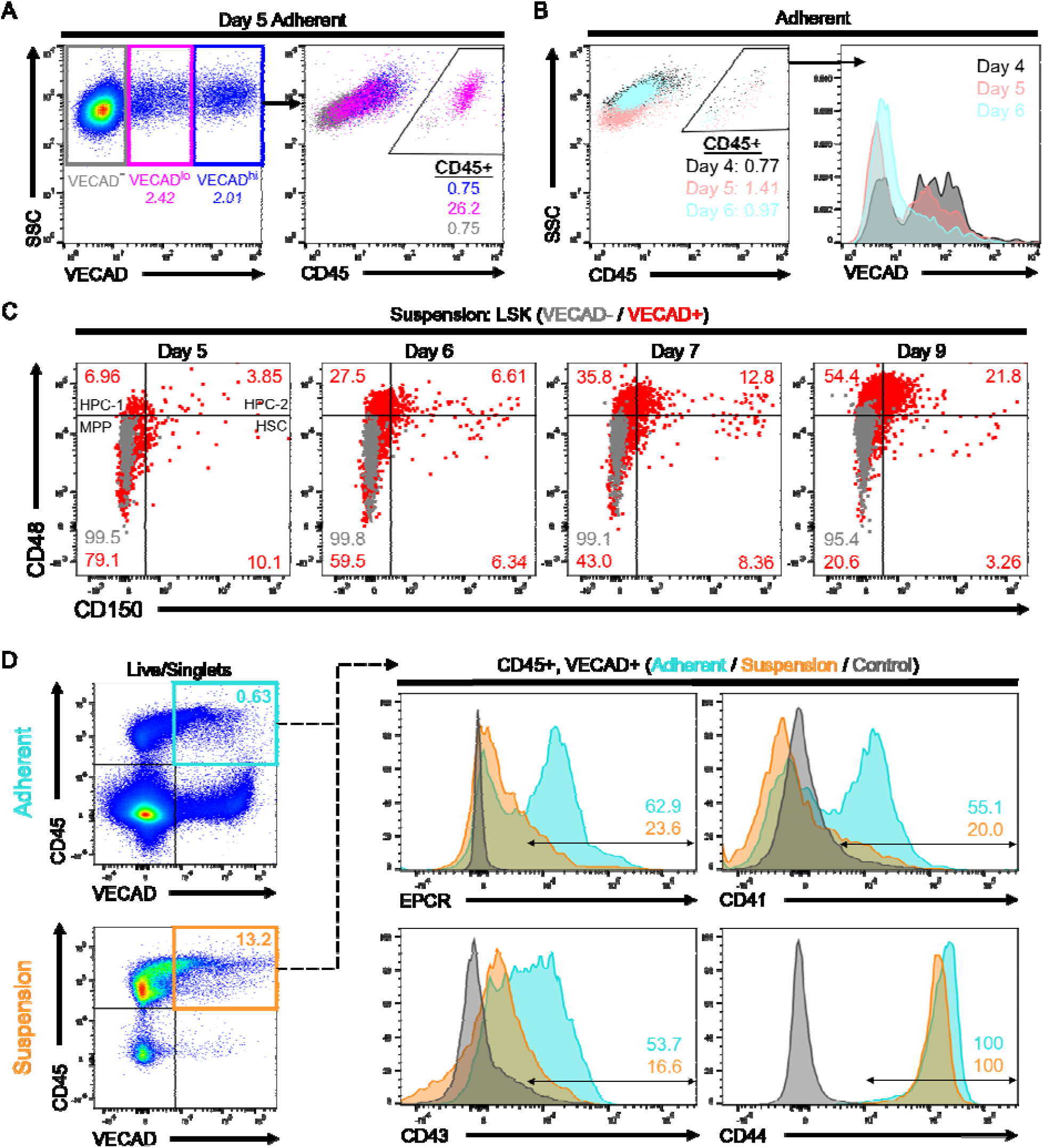
Murine fetal lung explants exhibit the dynamics of EHT. (A) Flow cytometric assessment of VECAD and CD45 expression on day 5 fetal lung explants. (B) CD45 and VECAD expression across days 4 to 6. VECAD expression is gradually lost as endothelial cells complete their transition to CD45^+^ hematopoietic cells, and this process diminishes across days 4-6 of culture. (C) Assessment of SLAM marker defined HSPC populations stratified by VECAD expression across multiple days in culture. (D) Assessment of pre-HSC and EHT markers on VECAD^+^/CD45^+^ defined progenitors and separated by adherent versus suspension cell populations.

Beyond VECAD, successive staging from a pro-HSC to a pre-HSC phenotype is marked by the expression of CD41 and CD43, respectively^40^. EPCR and CD44 have also been previously demonstrated as markers of pre-HSPCs^40–42^. Fetal lung derived VECAD^+^/CD45^+^ cells showed expression of CD41, CD43, CD44, and EPCR (**Figure 2D**). There was a notable reduction in CD41, CD43 and EPCR expression as these cells transitioned into a suspension state. Intensity of EPCR expression also decreased as VECAD^+^/CD45^-^ cells matured away from an endothelial signature (**Supplemental Figure 1B**). In contrast, CD41 expression transiently peaked at the VECAD^+^/CD45^+^ pre-HSPC stage. Under these experimental conditions, however, no notable changes in CD44 expression were observed regardless of physical and maturational state. We also observed significantly less expression of pre-HSC markers on fetal liver VECAD^+^/CD45^+^ cells suggesting that in contrast to those from the fetal lung, these cells are likely not pre-HSPCs (**Supplemental Figure 1A**).

### Human fetal lung explants undergo EHT to produce HSPCs

In an effort to understand if this phenomenon is conserved in humans, cells collected from human fetal lungs isolated from embryos aged between 19-24 weeks post-conception were cultured under the same conditions described above. Similar to murine fetal lung explants, human fetal lung explants initially developed an adherent layer and gave rise to suspension cells between days 7-10 (**Figure 3A**). Cytospins and flow cytometry of suspension cells showed progenitor-like cells and various differentiated cell types, including megakaryocytes, red blood cells, monocytes/macrophages and neutrophils (**Figure 3B, C**). Suspension cells assessed by Methocult assay also showed CFU capacity (**Figure 3D**). These data collectively suggest that human fetal lung explants are giving rise to HSPCs capable of multilineage differentiation.

**Figure 3.**
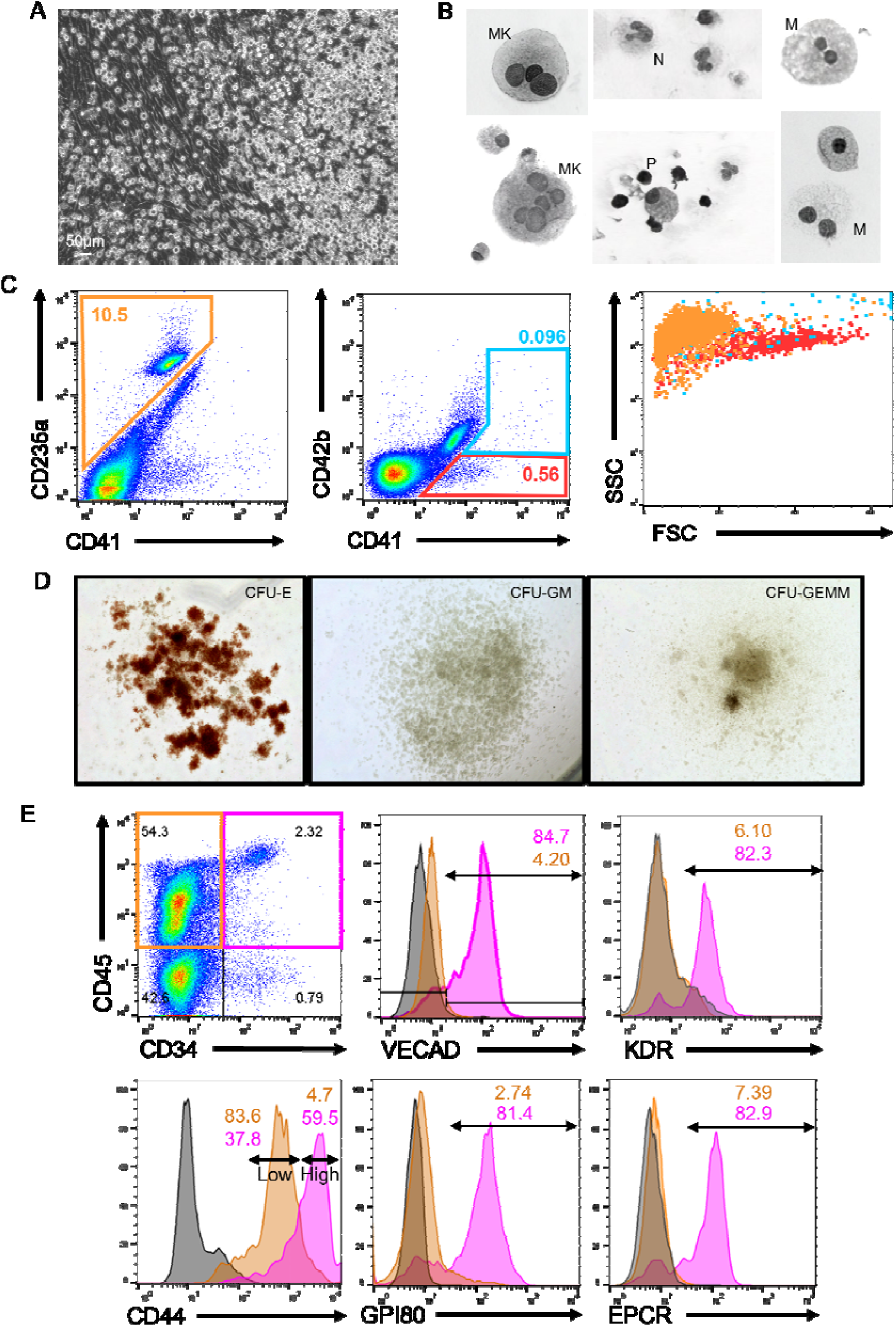
Human fetal lung explants undergo EHT to produce HSPCs. (A) Image of a human fetal lung explant culture showing a robust population of floating hematopoietic cells against a background of adherent cells. (B) Cytospin of human fetal lung suspension cells. (C) Flow cytometric assessment of differentiated erythrocyte and megakaryocyte populations. (D) Images of representative colonies from a CFU assay. (E) Assessment of pre-HSC and EHT markers on CD34^+^/CD45^+^ defined progenitors versus CD34^-^/CD45^+^ differentiated hematopoietic cells.

To assess whether these suspension cells were derived via EHT, flow cytometric assessment of endothelial and HSPC markers was performed on human fetal lung explants (**Figure 3E**). Human HSPCs were broadly defined by co-expression of the progenitor marker CD34 and the hematopoietic marker CD45. The majority of fetal lung derived CD34^+^/CD45^+^ progenitors co-expressed VECAD, which was downregulated as these cells differentiated and lost expression of CD34. Additionally, fetal lung derived HSPCs expressed the HSPC markers KDR, GPI80, CD44 and EPCR. Reduction in the expression of these pre-HSPC/HSPC markers coincided with the loss of CD34 expression. Notably, human fetal lung derived HSPCs were enriched with a distinct CD44-high population, which others have reported is a marker of type II pre-HSPCs found within the intra-aortic hematopoietic clusters of the AGM^41^.

Hematopoietic clusters are a common histological hallmark of EHT occurring within the AGM and can be found in other organs that exhibit EHT including the placenta, and the umbilical and vitelline arteries^8^. To investigate whether fetal lung EHT can occur in-vivo we performed immunofluorescent staining of fixed-frozen human fetal lung sections. Immunofluorescent analysis revealed distinct regions of colocalized cells co-expressing the broad hematopoietic marker CD45, the endothelial marker VE-Cadherin, and the pre-HSC marker CD41a (**Figure 4A**). Additional staining for E-Cadherin suggests that these regions of EHT are not localized within the developing lung epithelium (**Figure 4B**).

**Figure 4.**
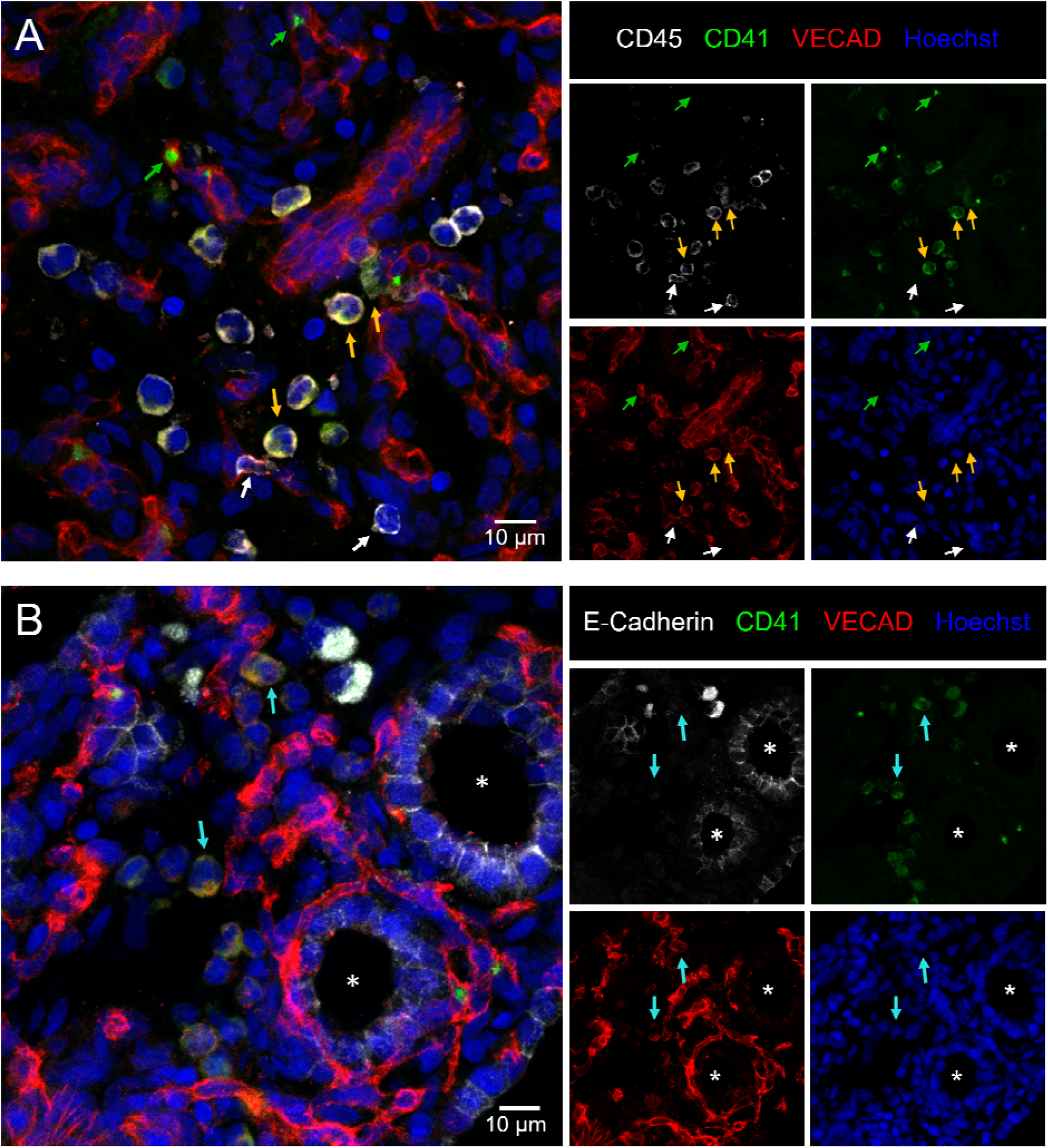
Immunofluorescent staining of in-situ fetal lung EHT. 12µm sections of a fixed frozen post-conception week 20 human fetal lung stained with the broad hematopoietic marker CD45, the EHT marker CD41, the endothelial marker VECAD, the epithelial cell marker E-Cadherin, and the nuclear stain Hoechst. (A) Cells co-expressing VECAD, CD41 and CD45 are highlighted with yellow arrows. Anucleate platelets single positive for CD41 are highlighted with green arrows. Differentiated hematopoietic cells only positive for CD45 are highlighted with white arrows. (B) Cells co-expressing VECAD and CD41 are highlighted with teal arrows, demonstrating they are not in developing E-Cadherin^+^ epithelial spaces, which are highlighted by a white asterisk.

### Single cell transcriptomic mapping of murine fetal lung EHT

To map the developmental trajectory of cells produced in the murine fetal lung explant model system and to illustrate the repertoire of cells involved in the process, Single cell RNA sequencing (scRNA-Seq) was performed. The optimal time window for EHT in our model was determined to be from days 4-6 with the peak of CD45^+^/VECAD^lo^ cells on day 5 (**Supplemental Figure** 2A). Since cells that have recently undergone EHT transiently retain expression of VECAD, adherent cells were collected into a single cell suspension and sorted for VECAD^+^ to enrich for both endothelial cells and cells undergoing transition (**Supplemental Figure 2B**). To ensure the capture of progenitor cells as well as differentiated cells that have downregulated VECAD post-EHT, suspension cells were collected for sequencing but left unsorted. The single cell capture and processing was performed using the Chromium 10x Genomics platform, and subsequently sequenced using the Illumina NextSeq 2000 platform. A schematic of the sequencing methodology as well as cell capture numbers and read depth can be found in **Supplemental Figure 2C, D**.

SPRING analysis revealed a trajectory of cells transitioning from an adherent to a suspension state through 3 distinct clusters: endothelial, transitional, and hematopoietic (**Figure 5A, B**). Supervised gene expression analyses of canonical endothelial markers showed robust expression and subsequent downregulation in the endothelial and transitional populations, respectively (**Figure 5C**). Analysis of genes previously described to be important during EHT, many of which were recently reviewed^3^, highlighted the expression of these key markers at a discrete stage of EHT. For example, *Notch1, Jag1, Dll4*, and *Sox17* were largely isolated to the endothelial cell stage. In contrast, *Yap1* and *Tead1* demonstrated sustained expression through the transitional cell stage. These results are in agreement with other reports demonstrating that *Notch1* and *Sox17* are critical in the early patterning of HECs, but their continued expression throughout EHT can be restrictive^43,44^. In contrast, Yap was previously shown to be critical in the maintenance of EHT, but not its initiation^15^. Lastly, expression of *Lyve1* was predominantly expressed during the endothelial stage, which suggests the production of a definitive wave of hematopoiesis^45(p1)^.

**Figure 5.**
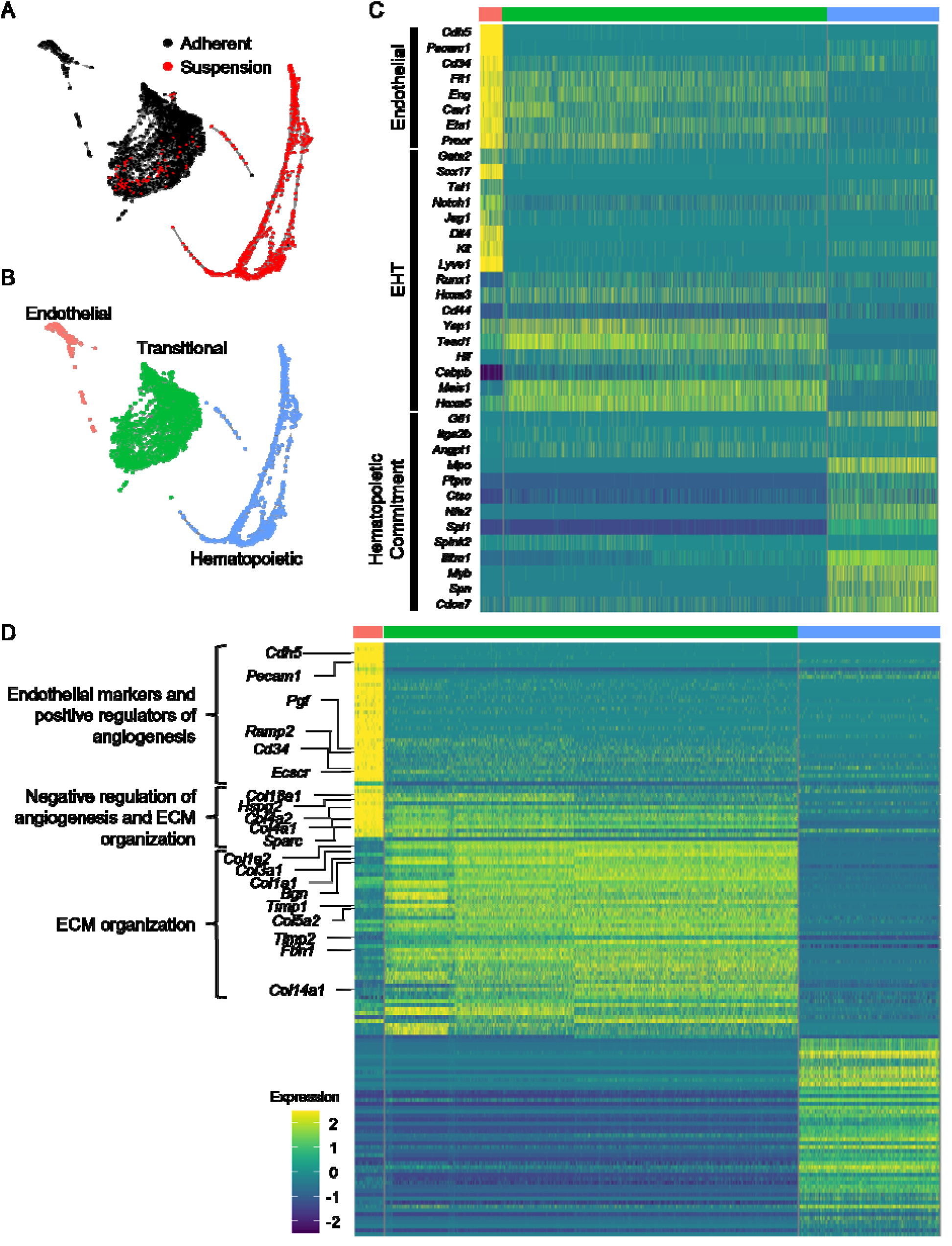
Single cell transcriptomic mapping of murine fetal lung EHT. (A, B) SPRING plot trajectory of EHT clusters. (C) Heatmap of supervised gene expression analysis of endothelial, EHT and hematopoietic commitment markers. (D) Heatmap of unsupervised analysis of the top 50 differentially expressed genes. Representative gene ontology analysis is highlighted here along with the associated genes enriched in these biological processes. Full DGE analysis can be found under supplemental data.

As cells transitioned to hematopoietic commitment, they began expressing key pre-HSPCs genes including *Cdca7, Myb, Gfi1* and *Spn* (CD43) (**Figure 5C**). Expression of *Cdca7* was previously demonstrated to be isolated to pre-HSC populations found within AGM intra-aortic hematopoietic clusters^46^. *Myb* is critical in HSC maintenance and proliferation^47^, *Gfi1* is critical in mediating the loss of an endothelial identity^48(p1)^, and *Spn* (CD43) is a known marker of early pre-HSCs^40,49^. Notably, expression of these genes was primarily isolated to early progenitor clusters and decreased along the trajectory of maturation (**Supplemental Figure 3A**), which is in agreement with other reports showing that their downregulation is necessary to allow for competent differentiation.

Complementary to these supervised analyses, unsupervised analysis of the top 50 differentially expressed genes showed that transitional cells retain the expression of genes involved in the downregulation of angiogenesis as they differentiate away from an endothelial fate (**Figure 5D, Supplemental Figure 3B, Supplemental**_**Data**). This coincided with the upregulation of genes critical in extracellular matrix (ECM) organization as these cells undergo a physical transition from an adherent to a suspension state. In agreement with these findings, others have demonstrated that the emergence of hematopoietic cells from HECs relies on the *Runx1* mediated upregulation of genes involved in ECM organization, cell adhesion and cell migration^50(p1)^. Many of the same genes, including *Runx1*, were upregulated predominantly within the transitional cluster (**Figure 5C, D**).

### Fetal Lung EHT is functionally reliant on canonical developmental pathways

To functionally assess the dependence of fetal lung EHT on Notch signaling, the Notch inhibitor, Compound E, was applied to explant cultures. Application of Compound E led to a significant reduction in VECAD^+^/CD45^+^ pre-HSPCs (**Figure 6B, C**). This is in line with other reports demonstrating that in-vitro Notch inhibition and in-vivo transgenic knockout of Notch1 specifically blocks EHT, but the proliferation, differentiation, and maintenance of HSPCs is conserved^43,51^.

**Figure 6.**
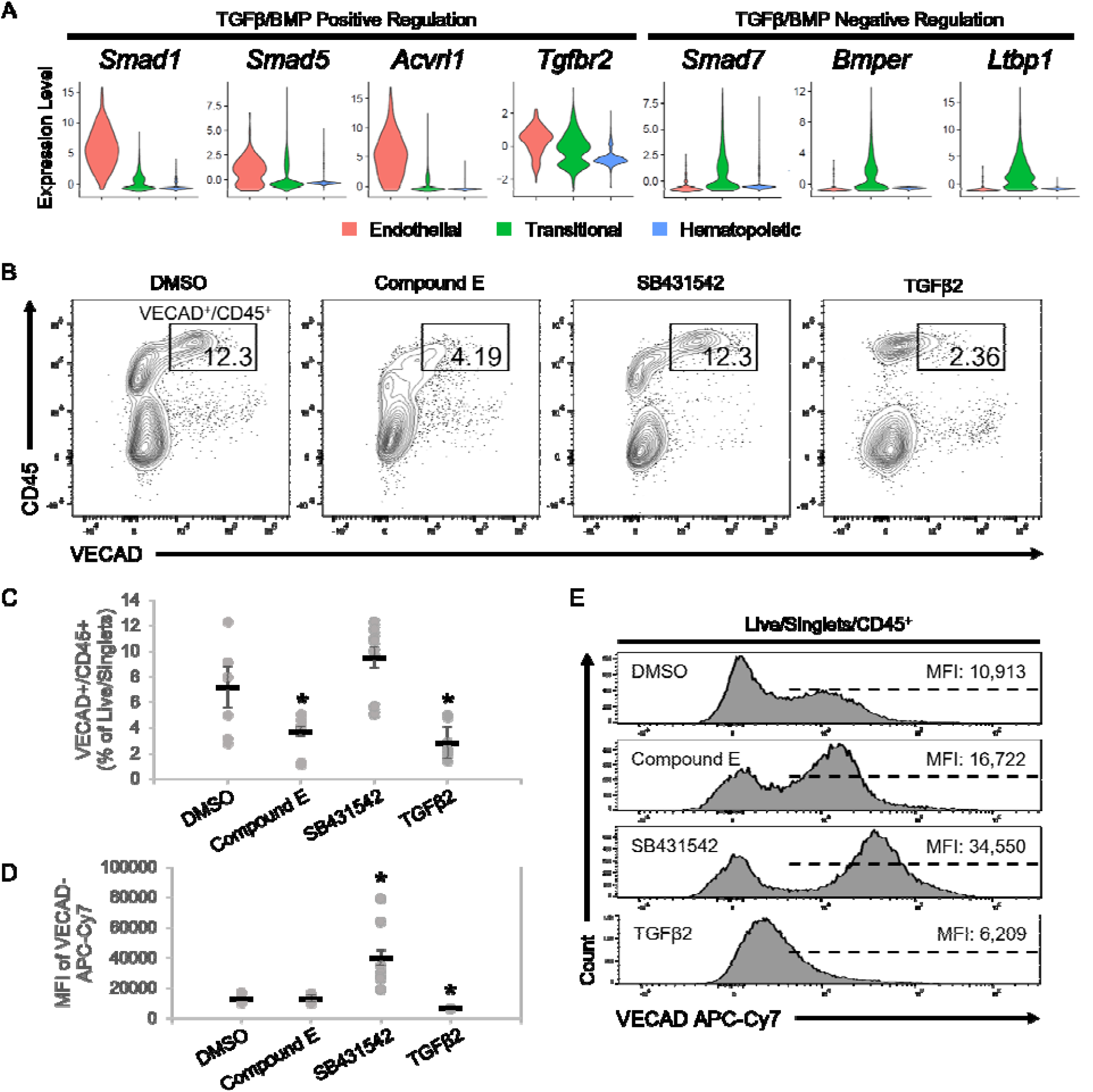
Fetal Lung EHT is functionally reliant on canonical developmental pathways. (A) Violin plots of expression of TGFβ/BMP pathways genes. (B) Representative flow cytometry plots demonstrating the change in CD45^+^/VECAD^+^ progenitors post-treatment with the Notch inhibitor Compound E, the TGFβ inhibitor SB431542, and recombinant TGFβ2. (C) Counts of CD45^+^/VECAD^+^ progenitors post-treatment. (D) Average MFI of VECAD expression post-treatment. (E) Representative histograms of VECAD expression post-treatment. Error bars represent standard deviation. *P < 0.01 compared to DMSO control.

In addition to the Notch family genes described previously, gene expression analysis demonstrated that fetal lung HECs exhibit the dynamic regulation of TGFβ/BMP pathways required for EHT. Similar to Notch, TGFβ/BMP pathways are important in the early patterning of HECs, but their subsequent inhibition is required for EHT^44,52–54^. Expression of key mediators of TGFβ/BMP signaling, including *Smad1, Smad5, Acvrl1* and *Tgfbr2*, peaked at the endothelial stage and was progressively downregulated throughout EHT (**Figure 6A**). Driving this downregulation was the concurrent upregulation of the TGFβ/BMP inhibitors *Smad7, Bmper* and *Ltbp1*.

To assess the dependence of TGFβ signaling in fetal lung EHT, the TGFβ receptor inhibitor, SB431542, and recombinant TGFβ2 were applied to explant cultures. Application of recombinant TGFβ2 led to a significant reduction in VECAD^+^/CD45^+^ pre-HSPCs (**Figure 6B, C**). In contrast, SB431542 did not significantly affect the production of VECAD^+^/CD45^+^ pre-HSPCs. However, SB431542 induced a significant increase in the intensity of VECAD expression, while TGFβ2 caused a reduction (**Figure 6D, E**). While others have shown that SB431542 treatment can enhance the production of HSPCs^44,55^, these findings suggest that there is already sufficient inhibition from the upregulation of TGFβ/BMP inhibitors discussed previously.

## DISCUSSION

Although not defined as a hematopoietic organ, the lung houses many resident blood cells that carry out both broad and tissue specific hematopoietic functions. Immune surveillance in the lung is carried out via unique resident immune cells, including dendritic cells, T-cells, B-cells, alveolar macrophages, interstitial macrophages, innate lymphoid cells and natural killer cells^11,56^. Supporting these classical immune cells are lung-resident megakaryocytes, which have a unique immune phenotype in addition to carrying out platelet biogenesis^57,58^. Lastly, the murine lung has been demonstrated to house HSPCs^12–14^. Complementary to these prior findings, we have demonstrated here that the murine and human fetal lungs are a source of functional HECs capable of giving rise to HSPCs. This suggests that the developing lung may have a more direct role in hematopoiesis than previously thought.

Although we have demonstrated the presence of fetal pulmonary HECs, the physiological significance of de-novo hematopoiesis occurring outside of the AGM and in the lung remains unclear. The potential need for the fetal lung to have an alternative source of blood may stem from the fact that most blood is shunted away from the fetal lung because oxygenation is provided via the mother. As a result, the developing murine lung only receives about 16% of total circulating blood^59^. The de-novo generation of blood in the fetal lung may act as an additional in-situ source of essential blood products for organogenesis. This is an important consideration when accounting for the role of macrophages and platelets in both organogenesis and angiogenesis^60–63^. Notably, the origins of many lung resident blood cells are not fully understood, but to date, the origins of resident macrophages are the most well studied.

Lung macrophages are derived from three distinct developmental waves originating from the yolk sac, fetal liver and BM^64^. Each wave persists into adulthood and occupies unique niches within the lung^65^. Recent work also suggests that such developmentally and phenotypically distinct subsets of macrophages may also extend to the heart, liver, kidney, and brain^66^. These findings demonstrate the complex diversity seen throughout hematopoietic development and how each phase individually makes significant physiological contributions. As such, the differentiation capacity of lung HECs requires further investigation to determine whether lung HECs contribute to the in-situ development of lung hematopoietic populations.

While often discussed as the precursor to HSCs, HECs also give rise to other blood progenitors. A subset of placental HECs lineage traced by *Hoxa13* preferentially form placental macrophages, termed Hofbauer cells, that remain solely in the placenta^67^. This finding also suggests that differentiation capacity and bias may already be predetermined at the HEC stage, which others have also proposed. THBS1 marks a subset of human embryonic stem cell derived endothelial cells that are biased towards megakaryopoiesis^68^. *Ly6a* versus *Tek* expression marks HECs with HSC versus EMP potential, respectively^69^. CXCR4 expression differentiates AGM derived HECs with HSC versus MPP potential^70^.

Others have described the presence of functional HSCs in E16 murine fetal lungs. However, LSK progenitors only make up on average about 0.02% of the total fetal lung cell isolate, compared to 0.75% in the adult BM^14^. This low number of HSPCs suggests that the role of fetal lung HECs may not contribute significantly to the progenitor cell pool and instead are skewed towards the immediate production of differentiated cell populations.

While more in-vivo work is required to determine the developmental trajectory and potency of lung HECs, the contributions of hematopoiesis that are not reliant on long-term HSCs are important to consider. Indeed, the majority of embryonic hematopoiesis is maintained by a pool of yolk sac derived erythro-myeloid progenitors (EMPs) that reside in the fetal liver^71,72^. EMPs give rise to the majority of tissue resident macrophages that persist into adulthood^73^, and embryonic erythrocyte production is sustained primarily from EMPs^74^. In addition to EMPs, transient HSCs that do not persist into adulthood have also been demonstrated to be involved in the establishment of the developing hematopoietic system. *Tie2*-Cre lineage tracing studies suggest that fetal HSCs that contribute to establishment of the developing hematopoietic system differentiate rapidly, which is in stark contrast to adult long term HSCs that are relatively quiescent^75^. Studies utilizing the *Flk2*-Cre tracer model (FlkSwitch) also demonstrate the existence of a transient population of HSCs with a lymphoid bias^76,77^. Although fulfilling the classic definition of an HSC based on their capacity for long-term multi-lineage reconstitution, these transient HSCs are only present from E10.5 until P14.

Notably, the fetal lung HECs described here were isolated from E17 mice, which is outside of the window of EHT within the AGM (E9.5-11.5). Work from Yvernogeau et al. also suggests that EHT can occur beyond this AGM window within the perinatal bone marrow^10^. They noted that perinatal sources of hematopoiesis occur at a time when HSC expansion in the fetal liver has stopped (∼E17) and the BM niche is still maturing in order to support the long-term residency of adult HSCs. Later stage EHT in the lung may thus aid in minimizing the need for long term HSC-dependent hematopoiesis during early development.

Niche interactions are another critical component of driving EHT^4,78^. Induction of key developmental pathways via secreted ligands is spatially organized within the AGM and influence the potency of resultant progenitors^20^. Replicating these conditions is thus critical in simulating EHT in an in-vitro setting. Most other employed methods of ex-vivo HEC cultures are reliant on the utilization of a supportive feeder cell layer, such as OP9 cells or immortalized AKT endothelial cells, to instruct EHT^51,79^. Notably, fetal lung explants undergo EHT without such feeders, which suggests that the lung independently has the cellular makeup to support EHT. However, further investigation is required to determine the cell-specific interactions of the in-vivo microenvironment where EHT would occur and how the lung niche instructs local HEC development.

Altogether, the findings described here demonstrate the existence of functionally validated HECs in the fetal lung, a population which is also conserved in humans. Given the diversity in HEC development and the significance of each distinct wave of hematopoiesis, the physiological influence of lung HEC development in-vivo needs further investigation. Expanding our overall understanding of HEC location and timing will also aid towards the goal of fully understanding the biological cues required for the development of functional HECs. Such findings can eventually be harnessed for the common goal of producing putative HSCs and functionally mature hematopoietic cells from pluripotent stem cells.

## LIMITATIONS OF STUDY

In this study, explant cultures were the primary methodology employed to investigate lung HECs. This ex-vivo methodology removes these cells from their microenvironment, thus limiting our understanding of these cells in their native in-vivo context. Of note, we do not believe we have induced the direct conversion of endothelial cells as direct conversion involves the forced overexpression of select transcription factors to competently induce transdifferentiation^3^. However, this still leaves many questions about whether these cells under steady state conditions in-vivo will naturally undergo EHT. Many tracing methods were considered to investigate these questions but also have significant limitations. While many lineage tracing models have been developed to trace either endothelial cell or HSPC populations individually, all models exhibit a significant amount of recombination events across both populations, especially during embryonic development^80^. As such, application of these methods would still not sufficiently answer such questions when considering the rarity of HECs and the existence of circulating HSPCs^81^, especially during the embryonic migration of HSPCs exiting the fetal liver. An inducible barcoding model was also considered^82^. However, others have previously noted that EHT is not the result of asymmetric cell division^37^. As such, a parent endothelial cell to trace the resultant hematopoietic progeny back to would not exist and negate the utility of such a method. Development of novel technologies to trace organ specific endothelial cell populations will greatly support these endeavors.

## ACKNOWLEDGEMENTS

This work was supported by the National Institutes of Health (NIH) Training Grant for Hematology (5T32HL7501-36) and the Predoctoral NRSA for MD/PhD Fellowships (1F30HL154552-01). The authors would like to thank the following for technical support: Brian R. Tilton of the Boston University Flow Cytometry Core Facility; Dr. Yuriy Alekseyev and Ashley LeClerc from the Boston University Microarray and Sequencing Resource (BUMSR) Core; Dr. Michael T. Kirber from the Boston University Cellular Imaging Core.

## AUTHORSHIP CONTRIBUTIONS

A.K.Y. performed experimental design, data collection, data analysis, interpretation of data and manuscript preparation. G.J.M. performed and supervised experimental design, interpretation of data and manuscript preparation. C.V.M. and J.L.V. performed the bioinformatic/computational analysis and supported the experimental design and manuscript preparation. A.C.B, K.V., T.W.D., A.B.Y., V.V., G.M. and A.B.B assisted with data collection and manuscript preparation.

## DISCLOSURES

The authors of this manuscript have no conflict of interest to disclose.

## Notes

### Competing Interest Statement

The authors have declared no competing interest.

## REFERENCES

1. Wu Y, Hirschi KK. Regulation of Hemogenic Endothelial Cell Development and Function. Annu Rev Physiol. 2021;83(1):17–37. doi:10.1146/annurev-physiol-021119-034352

2. Tavian M, Coulombel L, Luton D, Clemente HS, Dieterlen-Lievre F, Peault B. Aorta-Associated CD34+ Hematopoietic Cells in the Early Human Embryo. :6.

3. Lange L, Morgan M, Schambach A. The hemogenic endothelium: a critical source for the generation of PSC-derived hematopoietic stem and progenitor cells. Cell Mol Life Sci. Published online February 9, 2021. doi:10.1007/s00018-021-03777-y

4. Heck AM, Ishida T, Hadland B. Location, Location, Location: How Vascular Specialization Influences Hematopoietic Fates During Development. Front Cell Dev Biol. 2020;8:602617. doi:10.3389/fcell.2020.602617

5. Li W, Ferkowicz MJ, Johnson SA, Shelley WC, Yoder MC. Endothelial cells in the early murine yolk sac give rise to CD41-expressing hematopoietic cells. Stem Cells Dev. 2005;14(1):44–54. doi:10.1089/scd.2005.14.44

6. Chen MJ, Yokomizo T, Zeigler B, Dzierzak E, Speck NA. Runx1 is required for the endothelial to hematopoietic cell transition but not thereafter. Nature. 2009;457(7231):887–891. doi:10.1038/nature07619

7. Rhodes KE, Gekas C, Wang Y, et al. The Emergence of Hematopoietic Stem Cells is Initiated in the Placental Vasculature in the Absence of Circulation. Cell Stem Cell. 2008;2(3):252–263. doi:10.1016/j.stem.2008.01.001

8. Yzaguirre AD, Speck NA. Insights into blood cell formation from hemogenic endothelium in lesser-known anatomic sites. Developmental Dynamics. 2016;245(10):1011–1028. doi:10.1002/dvdy.24430

9. Li Z, Lan Y, He W, et al. Mouse Embryonic Head as a Site for Hematopoietic Stem Cell Development. Cell Stem Cell. 2012;11(5):663–675. doi:10.1016/j.stem.2012.07.004

10. Yvernogeau L, Gautier R, Petit L, et al. In vivo generation of haematopoietic stem/progenitor cells from bone marrow-derived haemogenic endothelium. Nature Cell Biology. 2019;21(11):1334–1345. doi:10.1038/s41556-019-0410-6

11. Ardain A, Marakalala MJ, Leslie A. Tissue-resident innate immunity in the lung. Immunology. 2020;159(3):245–256. doi:10.1111/imm.13143

12. Lefrançais E, Ortiz-Muñoz G, Caudrillier A, et al. The lung is a site of platelet biogenesis and a reservoir for haematopoietic progenitors. Nature. 2017;544(7648):105–109. doi:10.1038/nature21706

13. Summer R, Kotton DN, Liang S, Fitzsimmons K, Sun X, Fine A. Embryonic Lung Side Population Cells Are Hematopoietic and Vascular Precursors. Am J Respir Cell Mol Biol. 2005;33(1):32–40. doi:10.1165/rcmb.2005-0024OC

14. Hillel-Karniel C, Rosen C, Milman-Krentsis I, et al. Multi-lineage Lung Regeneration by Stem Cell Transplantation across Major Genetic Barriers. Cell Reports. 2020;30(3):807-819.e4. doi:10.1016/j.celrep.2019.12.058

15. Lundin V, Sugden WW, Theodore LN, et al. YAP Regulates Hematopoietic Stem Cell Formation in Response to the Biomechanical Forces of Blood Flow. Developmental Cell. 2020;52(4):446-460.e5. doi:10.1016/j.devcel.2020.01.006

16. Niblock MM, Perez A, Broitman S, Jacoby B, Aviv E, Gilkey S. In utero development of fetal breathing movements in C57BL6 mice. Respiratory Physiology & Neurobiology. 2020;271:103288. doi:10.1016/j.resp.2019.103288

17. Tang Z, Hu Y, Wang Z, et al. Mechanical Forces Program the Orientation of Cell Division during Airway Tube Morphogenesis. Developmental Cell. 2018;44(3):313-325.e5. doi:10.1016/j.devcel.2017.12.013

18. Li J, Wang Z, Chu Q, Jiang K, Li J, Tang N. The Strength of Mechanical Forces Determines the Differentiation of Alveolar Epithelial Cells. Developmental Cell. 2018;44(3):297-312.e5. doi:10.1016/j.devcel.2018.01.008

19. Canu G, Ruhrberg C. First blood: the endothelial origins of hematopoietic progenitors. Angiogenesis. 2021;24(2):199–211. doi:10.1007/s10456-021-09783-9

20. Souilhol C, Gonneau C, Lendinez JG, et al. Inductive interactions mediated by interplay of asymmetric signalling underlie development of adult haematopoietic stem cells. Nat Commun. 2016;7(1):10784. doi:10.1038/ncomms10784

21. Herriges M, Morrisey EE. Lung development: orchestrating the generation and regeneration of a complex organ. Development. 2014;141(3):502–513. doi:10.1242/dev.098186

22. Kaminow B, Yunusov D, Dobin A. STARsolo: Accurate, Fast and Versatile Mapping/Quantification of Single-Cell and Single-Nucleus RNA-Seq Data. Bioinformatics; 2021. doi:10.1101/2021.05.05.442755

23. Stuart T, Butler A, Hoffman P, et al. Comprehensive Integration of Single-Cell Data. Cell. 2019;177(7):1888-1902.e21. doi:10.1016/j.cell.2019.05.031

24. Hafemeister C, Satija R. Normalization and variance stabilization of single-cell RNA-seq data using regularized negative binomial regression. Genome Biology. 2019;20(1):296. doi:10.1186/s13059-019-1874-1

25. McInnes L, Healy J, Melville J. UMAP: Uniform Manifold Approximation and Projection for Dimension Reduction. arXiv:180203426 [cs, stat]. Published online December 6, 2018. Accessed June 5, 2020. http://arxiv.org/abs/1802.03426

26. Blondel VD, Guillaume JL, Lambiotte R, Lefebvre E. Fast unfolding of communities in large networks. J Stat Mech. 2008;2008(10):P10008. doi:10.1088/1742-5468/2008/10/P10008

27. Tirosh I, Izar B, Prakadan SM, et al. Dissecting the multicellular ecosystem of metastatic melanoma by single-cell RNA-seq. Science. 2016;352(6282):189–196. doi:10.1126/science.aad0501

28. Bourgon R, Gentleman R, Huber W. Independent filtering increases detection power for high-throughput experiments. PNAS. 2010;107(21):9546–9551. doi:10.1073/pnas.0914005107

29. Finak G, McDavid A, Yajima M, et al. MAST: a flexible statistical framework for assessing transcriptional changes and characterizing heterogeneity in single-cell RNA sequencing data. Genome Biology. 2015;16(1):278. doi:10.1186/s13059-015-0844-5

30. Soneson C, Robinson MD. Bias, robustness and scalability in single-cell differential expression analysis. Nat Methods. 2018;15(4):255–261. doi:10.1038/nmeth.4612

31. Kuleshov MV, Jones MR, Rouillard AD, et al. Enrichr: a comprehensive gene set enrichment analysis web server 2016 update. Nucleic Acids Res. 2016;44(W1):W90-97. doi:10.1093/nar/gkw377

32. Chen EY, Tan CM, Kou Y, et al. Enrichr: interactive and collaborative HTML5 gene list enrichment analysis tool. BMC Bioinformatics. 2013;14:128. doi:10.1186/1471-2105-14-128

33. Leung A, Zulick E, Skvir N, et al. Notch and Aryl Hydrocarbon Receptor Signaling Impact Definitive Hematopoiesis from Human Pluripotent Stem Cells: Definitive Hematopoiesis in Pluripotent Stem Cells. STEM CELLS. 2018;36(7):1004–1019. doi:10.1002/stem.2822

34. Yvernogeau L, Gautier R, Khoury H, et al. An in vitro model of hemogenic endothelium commitment and hematopoietic production. Development. 2016;143(8):1302–1312. doi:10.1242/dev.126714

35. Ohta R, Sugimura R, Niwa A, Saito MK. Hemogenic Endothelium Differentiation from Human Pluripotent Stem Cells in A Feeder- and Xeno-free Defined Condition. JoVE (Journal of Visualized Experiments). 2019;(148):e59823. doi:10.3791/59823

36. Boisset JC, van Cappellen W, Andrieu-Soler C, Galjart N, Dzierzak E, Robin C. In vivo imaging of haematopoietic cells emerging from the mouse aortic endothelium. Nature. 2010;464(7285):116–120. doi:10.1038/nature08764

37. Eilken HM, Nishikawa SI, Schroeder T. Continuous single-cell imaging of blood generation from haemogenic endothelium. Nature. 2009;457(7231):896–900. doi:10.1038/nature07760

38. Taoudi S, Gonneau C, Moore K, et al. Extensive Hematopoietic Stem Cell Generation in the AGM Region via Maturation of VE-Cadherin+CD45+ Pre-Definitive HSCs. Cell Stem Cell. 2008;3(1):99–108. doi:10.1016/j.stem.2008.06.004

39. Fantin A, Tacconi C, Villa E, Ceccacci E, Denti L, Ruhrberg C. KIT Is Required for Fetal Liver Hematopoiesis. Frontiers in Cell and Developmental Biology. 2021;9. Accessed February 12, 2022. https://www.frontiersin.org/article/10.3389/fcell.2021.648630

40. Ottersbach K. Endothelial-to-haematopoietic transition: an update on the process of making blood. Biochem Soc Trans. 2019;47(2):591–601. doi:10.1042/BST20180320

41. Oatley M, Bölükbasi ÖV, Svensson V, et al. Single-cell transcriptomics identifies CD44 as a marker and regulator of endothelial to haematopoietic transition. Nature Communications. 2020;11(1):1–18. doi:10.1038/s41467-019-14171-5

42. Fidanza A, Stumpf PS, Ramachandran P, et al. Single-cell analyses and machine learning define hematopoietic progenitor and HSC-like cells derived from human PSCs. Blood. 2020;136(25):2893–2904. doi:10.1182/blood.2020006229

43. Kumano K, Chiba S, Kunisato A, et al. Notch1 but Not Notch2 Is Essential for Generating Hematopoietic Stem Cells from Endothelial Cells. Immunity. 2003;18(5):699–711. doi:10.1016/S1074-7613(03)00117-1

44. Vargel Ö, Zhang Y, Kosim K, et al. Activation of the TGFβ pathway impairs endothelial to haematopoietic transition. Sci Rep. 2016;6(1):21518. doi:10.1038/srep21518

45. Lee LK, Ghorbanian Y, Wang W, et al. LYVE1 Marks the Divergence of Yolk Sac Definitive Hemogenic Endothelium from the Primitive Erythroid Lineage. Cell Rep. 2016;17(9):2286–2298. doi:10.1016/j.celrep.2016.10.080

46. Guiu J, Bergen DJM, De Pater E, et al. Identification of Cdca7 as a novel Notch transcriptional target involved in hematopoietic stem cell emergence. Journal of Experimental Medicine. 2014;211(12):2411–2423. doi:10.1084/jem.20131857

47. Greig KT, Carotta S, Nutt SL. Critical roles for c-Myb in hematopoietic progenitor cells. Seminars in Immunology. 2008;20(4):247–256. doi:10.1016/j.smim.2008.05.003

48. Lancrin C, Mazan M, Stefanska M, et al. GFI1 and GFI1B control the loss of endothelial identity of hemogenic endothelium during hematopoietic commitment. Blood. 2012;120(2):314–322. doi:10.1182/blood-2011-10-386094

49. Vodyanik MA, Thomson JA, Slukvin II. Leukosialin (CD43) defines hematopoietic progenitors in human embryonic stem cell differentiation cultures. Blood. 2006;108(6):2095–2105. doi:10.1182/blood-2006-02-003327

50. Lie-A-Ling M, Marinopoulou E, Li Y, et al. RUNX1 positively regulates a cell adhesion and migration program in murine hemogenic endothelium prior to blood emergence. Blood. 2014;124(11):e11–e20. doi:10.1182/blood-2014-04-572958

51. Hadland BK, Varnum-Finney B, Poulos MG, et al. Endothelium and NOTCH specify and amplify aorta-gonad-mesonephros–derived hematopoietic stem cells. J Clin Invest. 2015;125(5):2032–2045. doi:10.1172/JCI80137

52. Zhang C, Lv J, He Q, et al. Inhibition of endothelial ERK signalling by Smad1/5 is essential for haematopoietic stem cell emergence. Nat Commun. 2014;5(1):3431. doi:10.1038/ncomms4431

53. Monteiro R, Pinheiro P, Joseph N, et al. Transforming Growth Factor β Drives Hemogenic Endothelium Programming and the Transition to Hematopoietic Stem Cells. Dev Cell. 2016;38(4):358–370. doi:10.1016/j.devcel.2016.06.024

54. McGarvey AC, Rybtsov S, Souilhol C, et al. A molecular roadmap of the AGM region reveals BMPER as a novel regulator of HSC maturation. Journal of Experimental Medicine. 2017;214(12):3731–3751. doi:10.1084/jem.20162012

55. Wang C, Tang X, Sun X, et al. TGFβ inhibition enhances the generation of hematopoietic progenitors from human ES cell-derived hemogenic endothelial cells using a stepwise strategy. Cell Res. 2012;22(1):194–207. doi:10.1038/cr.2011.138

56. Barker KA, Etesami NS, Shenoy AT, et al. Lung-resident memory B cells protect against bacterial pneumonia. J Clin Invest. 2021;131(11). doi:10.1172/JCI141810

57. Pariser DN, Hilt ZT, Ture SK, et al. Lung megakaryocytes are immune modulatory cells. doi:10.1172/JCI137377

58. Yeung AK, Villacorta-Martin C, Hon S, Rock JR, Murphy GJ. Lung megakaryocytes display distinct transcriptional and phenotypic properties. Blood Advances. 2020;4(24):6204–6217. doi:10.1182/bloodadvances.2020002843

59. Zhou YQ, Cahill LS, Wong MD, Seed M, Macgowan CK, Sled JG. Assessment of flow distribution in the mouse fetal circulation at late gestation by high-frequency Doppler ultrasound. Physiological Genomics. 2014;46(16):602–614. doi:10.1152/physiolgenomics.00049.2014

60. Neo WH, Lie-A-Ling M, Fadlullah MZH, Lacaud G. Contributions of Embryonic HSC-Independent Hematopoiesis to Organogenesis and the Adult Hematopoietic System. Front Cell Dev Biol. 2021;9. doi:10.3389/fcell.2021.631699

61. Tsukiji N, Inoue O, Morimoto M, et al. Platelets play an essential role in murine lung development through Clec-2/podoplanin interaction. Blood. 2018;132(11):1167–1179. doi:10.1182/blood-2017-12-823369

62. Bertozzi CC, Schmaier AA, Mericko P, et al. Platelets regulate lymphatic vascular development through CLEC-2–SLP-76 signaling. Blood. 2010;116(4):661–670. doi:10.1182/blood-2010-02-270876

63. Battinelli EM. 24 -The Role of Platelets in Angiogenesis. In: Michelson AD, ed. Platelets (Fourth Edition). Academic Press; 2019:433–441. doi:10.1016/B978-0-12-813456-6.00024-2

64. Tan SYS, Krasnow MA. Developmental origin of lung macrophage diversity. Development. 2016;143(8):1318–1327. doi:10.1242/dev.129122

65. Chakarov S, Lim HY, Tan L, et al. Two distinct interstitial macrophage populations coexist across tissues in specific subtissular niches. Science. Published online March 15, 2019. doi:10.1126/science.aau0964

66. Dick SA, Wong A, Hamidzada H, et al. Three tissue resident macrophage subsets coexist across organs with conserved origins and life cycles. Science Immunology. Published online January 7, 2022. doi:10.1126/sciimmunol.abf7777

67. Liang G, Zhou C, Jiang X, et al. De novo generation of macrophage from placenta-derived hemogenic endothelium. Developmental Cell. 2021;56(14):2121-2133.e6. doi:10.1016/j.devcel.2021.06.005

68. Wang H, He J, Xu C, et al. Decoding Human Megakaryocyte Development. Cell Stem Cell. 2020;0(0). doi:10.1016/j.stem.2020.11.006

69. Chen MJ, Li Y, De Obaldia ME, et al. Erythroid/Myeloid Progenitors and Hematopoietic Stem Cells Originate from Distinct Populations of Endothelial Cells. Cell Stem Cell. 2011;9(6):541–552. doi:10.1016/j.stem.2011.10.003

70. Dignum T, Varnum-Finney B, Srivatsan SR, et al. Multipotent progenitors and hematopoietic stem cells arise independently from hemogenic endothelium in the mouse embryo. Cell Reports. 2021;36(11):109675. doi:10.1016/j.celrep.2021.109675

71. Dzierzak E, Bigas A. Blood Development: Hematopoietic Stem Cell Dependence and Independence. Cell Stem Cell. 2018;22(5):639–651. doi:10.1016/j.stem.2018.04.015

72. Frame JM, McGrath KE, Palis J. Erythro-Myeloid Progenitors: “definitive” hematopoiesis in the conceptus prior to the emergence of hematopoietic stem cells. Blood Cells Mol Dis. 2013;51(4):10.1016/j.bcmd.2013.09.006. doi:10.1016/j.bcmd.2013.09.006

73. Ghosn E, Yoshimoto M, Nakauchi H, Weissman IL, Herzenberg LA. Hematopoietic stem cell-independent hematopoiesis and the origins of innate-like B lymphocytes. Development. 2019;146(15):dev170571. doi:10.1242/dev.170571

74. Soares-da-Silva F, Freyer L, Elsaid R, et al. Yolk sac, but not hematopoietic stem cell–derived progenitors, sustain erythropoiesis throughout murine embryonic life. Journal of Experimental Medicine. 2021;218(e20201729). doi:10.1084/jem.20201729

75. Busch K, Klapproth K, Barile M, et al. Fundamental properties of unperturbed haematopoiesis from stem cells in vivo. Nature. 2015;518(7540):542–546. doi:10.1038/nature14242

76. Beaudin AE, Boyer SW, Perez-Cunningham J, et al. A transient developmental hematopoietic stem cell gives rise to innate-like B and T cells. Cell Stem Cell. 2016;19(6):768–783. doi:10.1016/j.stem.2016.08.013

77. Waas B, Maillard I. Fetal hematopoietic stem cells are making waves. Stem Cell Investig. 2017;4:25. doi:10.21037/sci.2017.03.06

78. Taoudi S, Medvinsky A. Functional identification of the hematopoietic stem cell niche in the ventral domain of the embryonic dorsal aorta. Proc Natl Acad Sci U S A. 2007;104(22):9399–9403. doi:10.1073/pnas.0700984104

79. Rybtsov S, Sobiesiak M, Taoudi S, et al. Hierarchical organization and early hematopoietic specification of the developing HSC lineage in the AGM region. Journal of Experimental Medicine. 2011;208(6):1305–1315. doi:10.1084/jem.20102419

80. Joseph C, Quach JM, Walkley CR, Lane SW, Lo Celso C, Purton LE. Deciphering Hematopoietic Stem Cells in Their Niches: A Critical Appraisal of Genetic Models, Lineage Tracing, and Imaging Strategies. Cell Stem Cell. 2013;13(5):520–533. doi:10.1016/j.stem.2013.10.010

81. Wright DE, Wagers AJ, Gulati AP, Johnson FL, Weissman IL. Physiological migration of hematopoietic stem and progenitor cells. Science. 2001;294(5548):1933–1936. doi:10.1126/science.1064081

82. Bowling S, Sritharan D, Osorio FG, et al. An Engineered CRISPR-Cas9 Mouse Line for Simultaneous Readout of Lineage Histories and Gene Expression Profiles in Single Cells. Cell. 2020;181(6):1410-1422.e27. doi:10.1016/j.cell.2020.04.048

