## Supplemental Figures and Tables for "DE-NOVO HEMATOPOIESIS FROM THE FETAL LUNG"

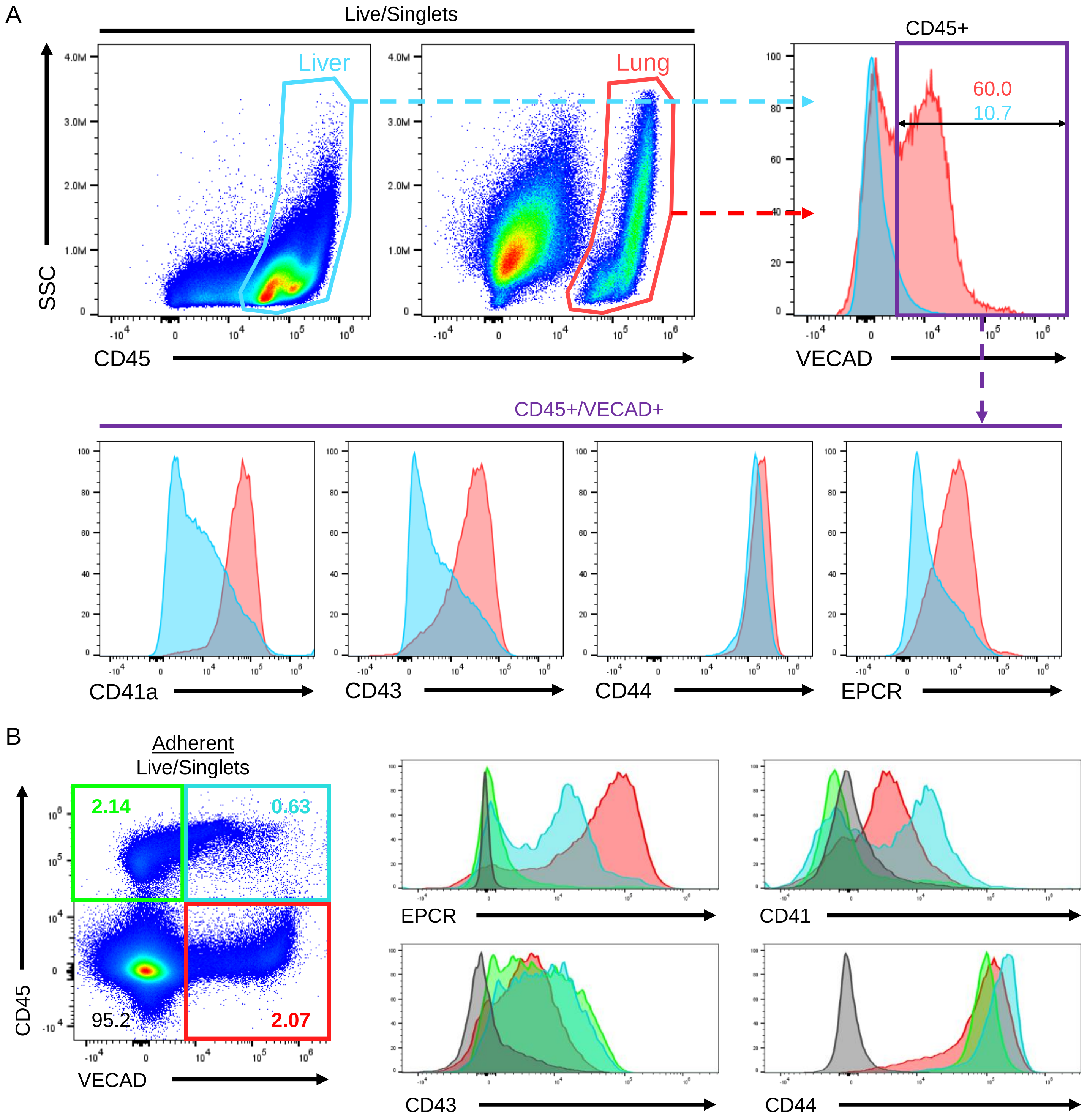


**Supplemental Figure 1. Assessment of EHT and pre-HSC markers on fetal liver and fetal lung explants.** (A) Comparison of EHT and pre-HSC markers between fetal liver and fetal lung explants. (B) Comparison of EHT and pre-HSC markers between populations differentiated by VECAD and CD45 expression.

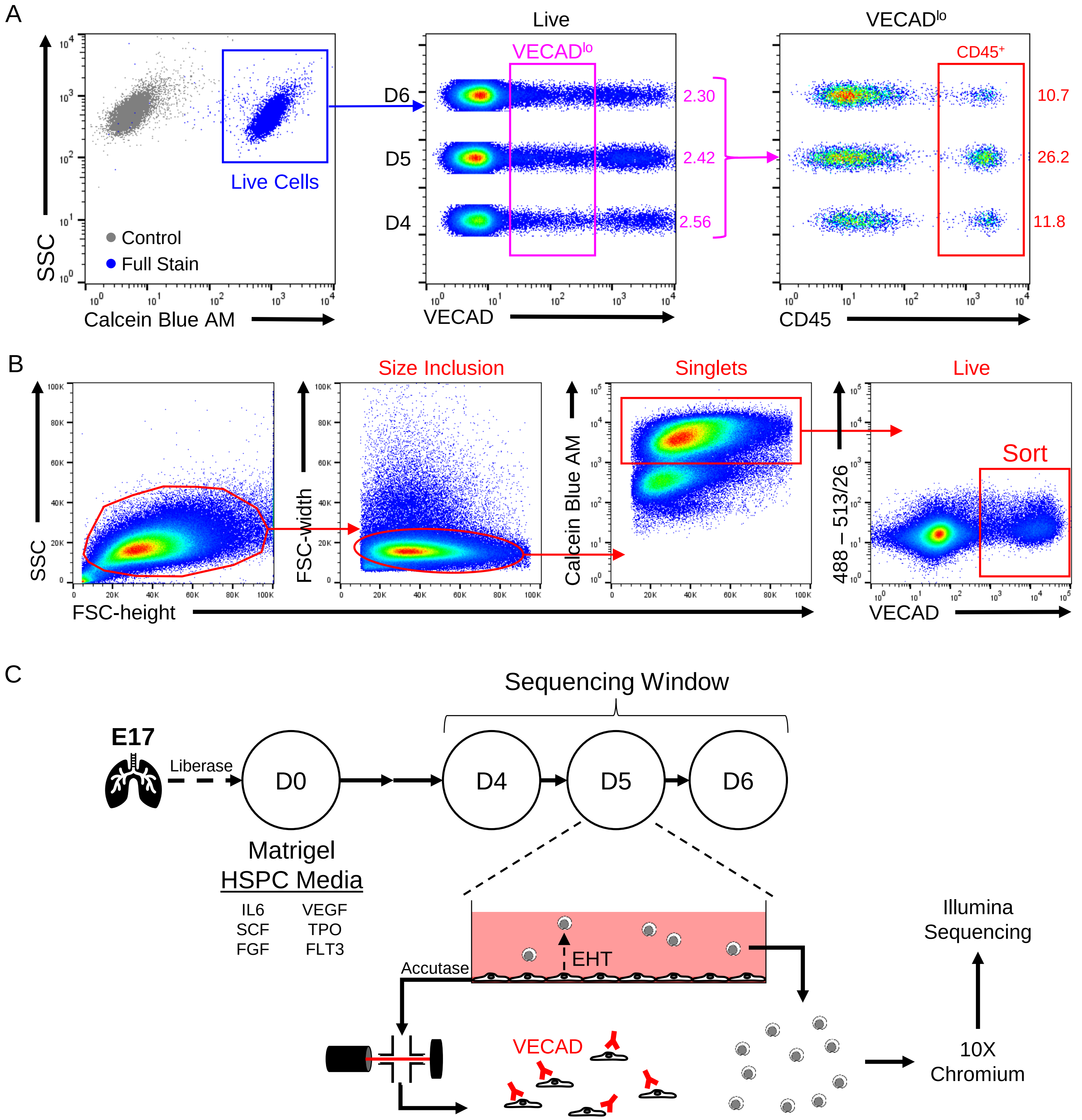


D

| Metric | Day 4 Suspension | Day 4 Adherent | Day 5 Suspension | Day 5 Adherent | Day 6 Suspension | Day 6 Adherent |
|---|---|---|---|---|---|---|
| Estimated Number of Cells | 2693 | 532 | 2497 | 1099 | 1398 | 1915 |
| Mean Reads per Cell | 16139 | 68754 | 17906 | 34019 | 39497 | 21106 |
| Median Genes per Cell | 2863 | 5681 | 2994 | 4098 | 3883 | 3162 |

**Supplemental Figure 2. Optimization of scRNA-Seq protocol and analysis.** (A) Comparison of the fraction of VECAD$^{lo}$ cells that are CD45$^{+}$. (B) Sorting strategy for scRNA-Seq of adherent cells from fetal lung explants. (C) Schematic of scRNA-Seq procedure. (D) Table showing baseline metrics of scRNA-Seq analysis.

A

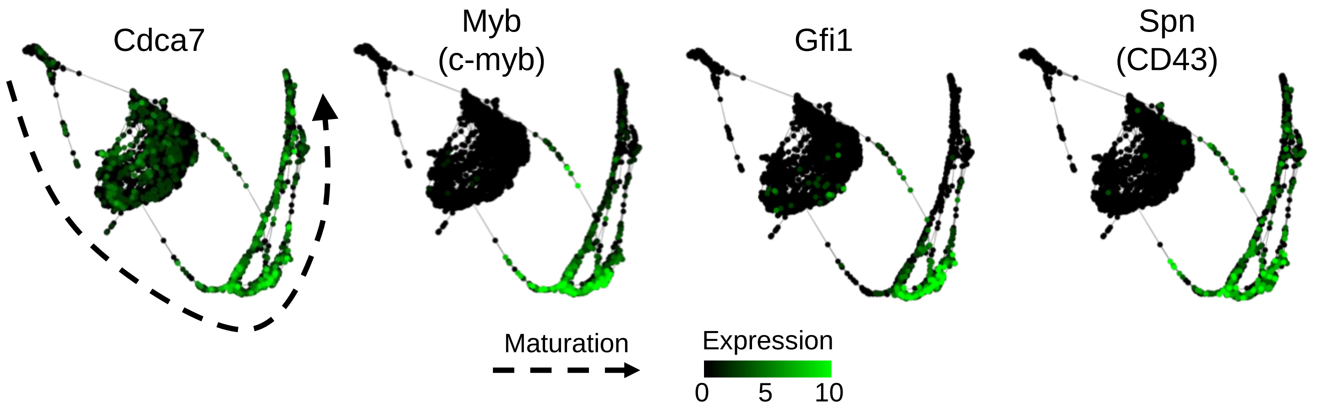


B

| Cluster | Index | Name | p-value |
|---|---|---|---|
| Endothelial | 1 | regulation of angiogenesis (GO:0045765) | 3.55E-08 |
| | 2 | apical junction assembly (GO:0043297) | 2.14E-06 |
| | 3 | bicellular tight junction assembly (GO:0070830) | 2.58E-05 |
| | 4 | blood vessel morphogenesis (GO:0048514) | 2.58E-05 |
| | 5 | tight junction assembly (GO:0120192) | 2.58E-05 |
| | 6 | glomerulus vasculature development (GO:0072012) | 2.58E-05 |
| | 7 | cell-cell junction assembly (GO:0007043) | 4.79E-05 |
| | 8 | maintenance of blood-brain barrier (GO:0035633) | 5.71E-05 |
| | 9 | extracellular structure organization (GO:0043062) | 5.71E-05 |
| | 10 | external encapsulating structure organization (GO:0045229) | 5.71E-05 |
| Transitional | 1 | extracellular matrix organization (GO:0030198) | 3.99E-05 |
| | 2 | skin morphogenesis (GO:0043589) | 0.000173 |
| | 3 | muscle contraction (GO:0006936) | 0.000173 |
| | 4 | extracellular structure organization (GO:0043062) | 0.000173 |
| | 5 | external encapsulating structure organization (GO:0045229) | 0.000173 |
| | 6 | skin development (GO:0043588) | 0.000222 |
| | 7 | supramolecular fiber organization (GO:0097435) | 2.69E-04 |
| | 8 | collagen fibril organization (GO:0030199) | 2.82E-04 |
| | 9 | regulation of membrane protein ectodomain proteolysis (GO:0051043) | 1.95E-03 |
| | 10 | negative regulation of dendritic cell differentiation (GO:2001199) | 4.07E-03 |
| Hematopoietic | 1 | neutrophil degranulation (GO:0043312) | 2.92E-13 |
| | 2 | neutrophil activation involved in immune response (GO:0002283) | 2.92E-13 |
| | 3 | neutrophil mediated immunity (GO:0002446) | 2.92E-13 |
| | 4 | defense response to bacterium (GO:0042742) | 2.61E-09 |
| | 5 | leukocyte aggregation (GO:0070486) | 0.000104 |
| | 6 | defense response to Gram-negative bacterium (GO:0050829) | 0.000112 |
| | 7 | neutrophil migration (GO:1990266) | 1.25E-04 |
| | 8 | innate immune response (GO:0045087) | 7.64E-04 |
| | 9 | regulation of phagocytosis (GO:0050764) | 8.85E-04 |
| | 10 | myeloid cell activation involved in immune response (GO:0002275) | 8.85E-04 |

**Supplemental Figure 3. SPRING and DGE analysis of fetal lung scRNA-Seq.** (A) SPRING plots showing the expression of key hematopoietic commitment markers. The dotted arrow represents the trajectory of maturation. (B) Extended list of Gene Ontology biological processes enrichment analysis. Full enrichment analysis can be found in Supplemental_Data.

| Marker | Species | Conjugate | Clone | Vendor | Catalog. # | Dilution |
|---|---|---|---|---|---|---|
| CD235a | Human | PE | HIR2 | BD Biosciences | 555570 | 1:1000 |
| CD34 | Human | BV421 | 581 | BD Biosciences | 562577 | 1:100 |
| CD41 | Human | BV421 | HIP8 | Biolegend | 303730 | 1:200 |
| CD41 | Human | Unconjugated | HIP8 | BD Biosciences | BDB555465 | 1:100 |
| CD42b | Human | APC | HIP1 | BD Biosciences | 551061 | 1:200 |
| CD45 | Human | APC | 2D1 | Biolegend | 368512 | 1:200 |
| CD45 | Human | PE-Cy7 | 2D1 | Biolegend | 368532 | 1:200 |
| EPCR (CD201) | Human | APC | RCR-401 | Biolegend | 351906 | 1:100 |
| GPI80 | Human | PE | 3H9 | MBL | D087-5 | 1:50 |
| KDR | Human | PE | 89106 | BD Biosciences | 560494 | 1:200 |
| KDR | Human | APC | 89106 | BD Biosciences | 560871 | 1:200 |
| Vascular Endothelial Cadherin (VECAD/CD144) | Human | AF647 | 55-7H1 | BD Biosciences | 561567 | 1:100 |
| Vascular Endothelial Cadherin (VECAD/CD144) | Human | PE | 55-7H1 | BD Biosciences | 561714 | 1:100 |
| Vascular Endothelial Cadherin (VECAD/CD144) | Human | Unconjugated | Polyclonal | R&D Systems | AF938 | 1:100 |
| CD44 | Human/Murine | BV711 | IM7 | Biolegend | 103057 | 1:800 |
| CD45 | Human/Murine | Unconjugated | Polyclonal | Invitrogen | PIPA587427 | 1:100 |
| Epithelial Cadherin (E-Cadherin) | Human/Murine | Unconjugated | DECMA-1 | Santa Cruz Biotechnology | sc-59778 | 1:50 |
| CD150 | Murine | BV711 | TC15-12F12.2 | Biolegend | 115941 | 1:400 |
| CD41 | Murine | BV421 | MWReg30 | Biolegend | 133911 | 1:400 |
| CD43 | Murine | FITC | S11 | Biolegend | 143204 | 1:100 |
| CD45 | Murine | BUV395 | 30-F11 | BD Biosciences | 564279 | 1:200 |
| CD48 | Murine | BV605 | HM48-1 | Biolegend | 103441 | 1:100 |
| EPCR | Murine | APC | eBio1560 | eBioscience | 50-151-23 | 1:200 |
| Kit | Murine | BV510 | ACK2 | Biolegend | 135119 | 1:400 |
| Kit | Murine | APC | 2B8 | Biolegend | 105812 | 1:200 |
| Lineage Cocktail | Murine | FITC | Cocktail | Invitrogen | 22777072 | 1:5 |
| Sca-1 (Ly-6A/E) | Murine | PE | D7 | eBioscience | 50-110-49 | 1:200 |
| Vascular Endothelial Cadherin (VECAD/CD144) | Murine | Biotin | BV13 | Biolegend | 138008 | 1:200 |
| Streptavidin | N/A | APC-Cy7 | N/A | Biolegend | 405208 | 1:200 |
| Calcein Blue AM | Viability Dye | N/A | N/A | Invitrogen | C1429 | 1:1000 |
| Live/Dead Fixable Blue | Viability Dye | N/A | N/A | Invitrogen | L34961 | 1:800 |

**Supplemental Table 1. Antibodies used for flow cytometry and immunofluorescent microscopy.**
